# On the concept of biological function, junk DNA and the gospels of ENCODE and Graur *et al.*

**DOI:** 10.1101/000588

**Authors:** Claudiu I. Bandea

## Abstract

In a recent article entitled “*On the immortality of television sets: “function” in the human genome according to the evolution-free gospel of ENCODE*”, Graur *et al*. dismantle ENCODE’s evidence and conclusion that 80% of the human genome is functional. However, the article by Graur *et al.* contains assumptions and statements that are questionable. Primarily, the authors limit their evaluation of DNA’s biological functions to informational roles, sidestepping putative non-informational functions. Here, I bring forward an old hypothesis on the evolution of genome size and on the role of so called ‘junk DNA’ (jDNA), which might explain C-value enigma. According to this hypothesis, the jDNA functions as a defense mechanism against insertion mutagenesis by endogenous and exogenous inserting elements such as retroviruses, thereby protecting informational DNA sequences from inactivation or alteration of their expression. Notably, this model couples the mechanisms and the selective forces responsible for the origin of jDNA with its putative protective biological function, which represents a classic example of ‘*fighting fire with fire*.’ One of the key tenets of this theory is that in humans and many other species, jDNAs serves as a protective mechanism against insertional oncogenic transformation. As an adaptive defense mechanism, the amount of protective DNA varies from one species to another based on the rate of its origin, insertional mutagenesis activity, and evolutionary constraints on genome size.

---

In a recent article entitled *“On the immortality of television sets: “function” in the human genome according to the evolution-free gospel of ENCODE”* [1], Graur *et al*. dismantle ENCODE’s evidence and conclusion that 80% of the human genome is functional [2], which would render the traditional concept of junk DNA (jDNA) as non-functional or parasitic DNA obsolete. I agree with many assertions made by the authors about the misleading macro-interpretation of data and poor rationale behind ENCODE’s conclusion; however, their article contains assumptions and statements that are questionable.

According to Graur *et al*., the ENCODE’s conclusion was based on a faulty, non-evolutionary definition of biological function. To make their point, the authors discuss at length the concept of biological function, differentiating between a “selected effect”, which is a historical and evolutionary perspective on function, and a “causal role”, an ahistorical and non-evolutionary perspective, and state that: “Armed with the proper concept of function, one can derive expectations concerning the rates and patterns of evolution of functional and nonfunctional parts of the genome. The surest indicator of the existence of a genomic function is that losing it has some phenotypic consequence for the organism”. Consistent with this rationale, the authors point to the results of comparative genomic studies, which show that, based on sequence conservation criteria, the total fraction of the human genome that is “certain to be functional” is approximately 9%, indeed, a far cry from ENCODE’s 80% figure.

However, Graur *et al*. limit their evaluation of genomic DNA’s biological functions to its informational roles (iDNA), which are based on sequence specificity. Although the authors recognize, apparently as an afterthought, that: “It has been pointed to us that…some parts of the genome may be functional but not under constraint with respect to nucleotide composition”, they sidestep the significance of non-informational roles for DNA (niDNA). True, only minute amounts of the genome have been attributed definite non-informational functions, some of which were mentioned in the article [1], but several, have been developed to explain the evolution and the putative biological functions of niDNA. For example, the ‘nucleo-skeletal’ [3] and ‘nucleotypic’ [4] functions, which “describe genome size variation as the outcome of selection via intermediate of cell size” [5], have been discussed in dozens of publications during the last few decades. Because the ‘nucleo-skeletal’ and ‘nucleotypic’ functions attributed to jDNA might represent adaptations of the host to the presence of jDNA (see below) rather than genuine biological functions, I will not elaborate on them here. Instead, in order to evaluate putative non-informational biological functions for jDNA, I bring forward an old hypothesis on the evolution of genome size and the function of jDNA, which might explain the C-value enigma [6; see Suplementary material].

According to this hypothesis, the so called jDNA functions as a defense mechanism against insertional mutagenesis by endogenous and exogenous mobile elements, thereby protecting iDNA sequences from inactivation or alteration of their expression. Notably, this model couples the mechanisms and the selective forces responsible for the origin of jDNA from inserting elements (see below) with its putative biological function as a defense mechanism against insertional mutagenesis by endogenous and exogenous inserting elements. Indeed, similar to the CRISPR system, in which viral sequences have been co-opted as an adaptive antiviral defense system [7], the defense mechanism provided by jDNA is a classic case of ‘*fighting fire with fire*’.

The rationale for this model was based on a broad evolutionary framework addressing two critical issues: (i) the mechanisms and selective forces leading to the origin of jDNA sequences, and (ii) the mechanisms and selective forces controlling the location and the quantity of jDNA sequences within the genome:

## (i) Origin of jDNA sequences

Approximately half of the human genome consists of recognizable endogenous viruses and transposable elements, and much of the remaining jDNA is composed of remnants of these elements. Therefore, the mechanisms and selective forces behind the genesis of jDNA sequences are associated primarily with the inserting elements, not with the host; however, it is important to note that all genomic sequences, including jDNA, also undergo duplications or amplifications during replication, recombination and chromosomal segregation (e.g. polyploidizasion), which represent additional mutational events leading to an increase in genome size.

Similar to all mutational events, occasionally, some of the inserting DNA sequences are beneficial for the host and, therefore, undergo positive selection. If the insertions cause deleterious effects, such as disrupting the coding and regulatory regions of the genome (see next section), they undergo negative selection. Even if not disruptive, in long term, the inserted DNA sequences experience negative selection because of associated genomic maintenance costs. In most organisms, including Bacteria, Archaea and many single-cell eukaryotic organisms, the purifying selection against non-functional or parasitic DNA is relatively strong. However, in many eukaryal organisms, including most multicellular species, the costs for maintaining these sequences are small compared to those associated with other organismal features; therefore, the purifying selection against the accumulation of parasitic genomic DNA is relatively weak, at least up to a certain quantity.

Overall, if the rate of its production is higher than that of its deletion, non-functional, or parasitic DNA can accumulate in the host’s genome. In this scenario, in order to accommodate for the genomic presence of large quantities of parasitic DNA, the hosts will adapt by adjusting some of their features, such as nucleotide metabolism or nuclear volume and cell size (6; see statement about ‘nucleo-skeletal’ and ‘nucleotypic’ theories above). These metabolic and physiological adaptations by the host to the presence of jDNA in their genome are highly relevant because experimental deletion of large quantities of jDNA (in order to prove or disprove its function) might have negative phenotypic consequences even if jDNA is non-functional, which questions the approach suggested by Graur *et al.*’ and others to define biological functions.

## (ii) Evolutionary constraints on the location and the amount of genomic jDNA

Key to exploring the hosts’ evolutionary constraints on the location and the amount of genomic jDNA, as well as it’s putative protective function, is the evolution of defense mechanisms in form of preferred or specific genomic sites for the integration of inserting elements in microbial organisms such as Bacteria, which have little jDNA. The evolution of these protective mechanisms is strong evidence for the selective pressure against insertional mutagenesis in these single-cell organisms. This selection pressure, however, take a new dimension in multicellular organisms, in which insertional mutagenesis occurs not only in the germline, but also in the somatic cells. Although the number of somatic insertional mutations during the course of the reproductive life span of multicellular organisms is enormous, because of the high turnover of cells in many tissues, most insertional mutations, including those causing cellular death, have a limited negative impact on the organism. The major problem is with the insertional mutagenesis that causes uncontrolled proliferation of cells, which can lead to neoplastic transformations, or cancer.

In humans, for example, given the enormous number of somatic cells and their high turnover rate during reproductive span, without protective mechanisms, the number of insertion events, especially those associated with exogenous viruses such as retroviruses, that could lead to cancer would be evolutionarily drowning. A dramatic example of the problems associated with insertional oncogenic transformation is from the highly promising biomedical field of gene therapy using viral vectors, which has been devastated by high prevalence of cancer in treated patients [8-12]. It is relevant to mention also that insertional transformation has been one of the main and most effective approaches for identifying and mapping genes and regulatory elements implicated in cancer [13-15], which points to the tremendous selection pressure imposed by cancer-inducing insertion mutagenesis in multicellular organisms. The protective function of jDNA against cancer development can be easily addressed, both analytically and experimentally; for example, transgenic mice carrying genomic DNA sequences homologous to infectious retroviruses, such as murine leukemia viruses (MuLV), might be more resistant to cancer induced by experimental retroviral (e.g. MuLV) infections as compared to controls.

Another strong line of evidence for the extraordinary selective pressure imposed by insertional mutagenesis and for the selective forces controlling the site of integration is the evolution of spliceosome, one of the most complex eukaryal macromolecular machineries [16]. The current prevalent view is that splicing machinery originated from group-II self-splicing introns as a defense system against insertional mutagenesis of iDNA [17-19]. Indeed, the evolution of introns and spliceosomes allows the insertion and accumulation of jDNA sequences within the transcribed regions of the genome, which often represent preferred regions for the integration of viral elements.

Given the strong negative impact of insertional mutagenesis, the evolution of protective mechanisms in form of specific integration sites and jDNA makes sense. This strong selective pressure led also to the evolution of additional defense molecular mechanisms, such as the Piwi-interacting RNAs (piRNAs), the largest class of small non-coding RNA molecules expressed in animal cells, and the RNA interference system (RNAi), which have evolved as specialized arms of the immune system defending against transposable elements and viruses [20, 21]. Once evolutionarily fixed, some of the components associated with these defensive systems, including some of the jDNA sequences, have been co-opted for other biological functions, such as gene regulation. It is relevant to emphasize at this point that not all phenotypes or biological functions are equally ‘evident’, or follow Graur *et al*.’s definition. For example, the human immune system contains hundreds if not thousands of components and eliminating some of them might not have immediate phenotypic effects, although it might have long term negative evolutionary consequences. Moreover, some of the genomic sequences coding for the conventional immune components, such as those implicated in the production of the repertoires of antibodies, T-cell receptors and MHC antigens, have been specifically selected against sequence conservation and, similar to the putative protective role of jDNA sequences, their protective function (i.e. phenotype) is fully attainable only as a ‘group activity’.

Assuming a random integration site, the protection level of jDNA would be directly proportional to its amount; nevertheless, other mechanisms that target the inserting elements into jDNA, such as preferential sites of integration and homologous recombination, could dramatically increase the protective role of jDNA (see, for example, the site specific integration of chromoviruses, an ancient and widespread lineage of Ty3-gypsy retrotransposons [22]). One of the most interesting tenets associated with the model discussed here is that, similar to the CRISPR system (see above), the jDNA defense system has a build-in adaptive feature, in the sense that an increase in the insertional activity would increase in the amount of jDNA, which would increase its potential protective function.

Whether functional or non-functional, genomic DNA sequences undergo deletion, a process that usually occurs during replication and recombination events. Evidently, mutational events consisting of deletions of functional sequences would enter negative selection, whereas deletions of parasitic DNA enter positive selection. According to Graur *et al*., though, “In humans, there seems to be no selection against excess genomic baggage”. However, non-functional or parasitic DNA is under purifying selection in all organisms, although less in some than in others, and there is eloquent evidence on evolutionary constraints on very large genomes [23-25]; in other words “without selection against excess genomic baggage” the human genome might be much larger.

Perhaps one of the most revealing examples of genome size evolution is found in cryptophytes and chlorarachniophytes, which contain 4 evolutionary distinct genomes [26]. The algal endosymbionts of these species have a small nucleus (called nucleomorph) with a genome ranging from ∼330 to 1,030 kilobase pairs, which is within the range of viral genomes. Compared to their ancestors, the genomes of these endosymbiotic algae have been reduced more than 200 fold. Remarkably, the number of their introns and their size have undergone drastic reductions, culminating with elimination of all introns and most, but not all, components of the spliceosomal machinery in at least in one species, *H. andersenii* [27]. The evolution of these remarkable endosymbiotic algae support the notion that deletion mutagenesis and the selection forces for eliminating jDNA, including introns, can be highly efficient in eukaryal genomes. In the context of the model discussed here, it is important to emphasize that, unlike the genome of their free living ancestors or that of most other eukaryotic cells, the genome of these endosymbionts is separated from host cytoplasm by several membranes (the nuclear envelope and the cellular and phagosomal membranes [26]), which constitute an effective ‘physical’ barrier and defensive system against exogenous inserting viral elements [18]. In the absence of newly introduced viral elements, the selective pressure associated with insertional mutagenesis had diminished, which led to the elimination of introns and of most jDNA, which are no longer needed as protective mechanisms; interestingly, the presence of membranes and the lack of mobile elements in these endosymbionts might also be responsible for the lack of transfer of their genes to the host genome and, therefore, for their evolutionary survival as nucleomorphs [28].

One of the most bizarre, but highly intriguing genome defense systems against invading inserting elements is found in ciliates, a highly diverse group of protozoans [29]. These organisms have two genomes: a germ-line, diploid genome, which is transcriptionally silent and carries tens of thousands of mobile elements, and a transcriptionally active polyploid genome, which originates from the germ-line genome by programmed DNA rearrangement and elimination of mobile elements. In some groups of ciliates, such as *Oxytricha*, over 90% of the germ-line genome is composed of jDNA, which is eliminated during the programmed DNA deletion. Apparently, in these single-cell organisms, maintaining the jDNA as a defense system against insertional mutagenesis in the germ-line genome was under very strong positive selection.

Among the most cited cases pertaining to the C-value enigma and the idea of jDNA as non-functional DNA is the genome of some amoeba species, which have the largest genome of all studied organisms. Along with other unusually large genomes, the amoeba genome are considered very strong evidence that most jDNA sequences do not have a biological function [1]. Indeed, the proponents of functional jDNA (e.g. ENCODE) have yet to explain the gigantic amoeba genome in context of their theories; nor have they addressed their perspective in context of the c-value enigma. Within the framework of the model presented here, the enormous genome of amoeba represent an extreme jDNA-based protective mechanism that has evolved in association with their huge appetite [30] for ingesting and hosting a myriad of microorganisms and their inserting elements.

## Perspective

Whether jDNA has been evolutionary maintained simply because of a mutational imbalance, favoring amplification of parasitic DNA versus deletion, or because jDNA is under host positive selection (whatever this selection might be), the protective function of jDNA in humans and other eukaryal organisms against insertional mutagenesis by endogenous and exogenous mobile genetic elements, such as retroviruses, is a *bona fide* fact. The abundant genomic presence of inserting elements and the evolution of several highly complex molecular defense mechanisms against insertional mutagenesis, including the splicing machinery, RNAi, programmed DNA deletion, methylation and repeat-induced point mutation defense system [31], testify for the extraordinary selective pressure imposed on the host genome by the endogenous and exogenous viral elements. In light of this selection pressure and of the fact that jDNA does provide protection against isertional mutagenesis, it is highly plausible that jDNA has been under positive selection for this critical biological function.

Unlike the selective forces acting upon the site of integration of jDNA sequences, which are strong and self-evident, those controlling the amount jDNA in most multicellular organisms might be weaker, multidimensional and more difficult to define. Nevertheless, according to the model discussed here, the amount of protective DNA as an adaptive defense mechanism varies from one species to another based on the rate of its origin, insertional mutagenesis activity, and evolutionary constraints on genome size.

In another recent critique of the ENODE’s conclusion, which also discusses in detail the concept of biological function and the C-value paradox, Ford Doolittle predicts that by building an informed theoretical framework on genome evolution “Much that we now call junk could then become functional” [32]. I think, we can reasonably state that, similar to hundreds of components of the immune system acting at the molecular, cellular, or organismal level, jDNA represent a broad and efficient molecular protective system against insertional mutagenesis and, therefore, it plays a significant biological role.

One of the main goals of the ENCODE project was to provide genomic insights into human health and disease, such as cancer. So far, this heavily funded project has yet to have a significant impact on our knowledge about cancer and other diseases. In contrast, one of the key tenets of the model discussed here is that jDNAs serve as a protective mechanism against insertional oncogenic transformation in humans and other multicellular species. Given the potential significance and implications of this model for one of the most devastating human diseases, as well as for understanding the evolution of genome size and resolving the long-standing C-value enigma, it would make sense to fully evaluate it, both theoretically and experimentally.

## Acknowledgments

I thank Dan Graur for his feedback.

